# Developmental depression-facilitation shift controls excitation-inhibition balance

**DOI:** 10.1101/2021.02.23.431593

**Authors:** David W. Jia, Rui Ponte Costa, Tim P. Vogels

## Abstract

Changes in the short-term dynamics of excitatory synapses over development have been observed throughout cortex, but their purpose and consequences remain unclear. Here, we propose that developmental changes in synaptic dynamics buffer the effect of slow inhibitory long-term plasticity, allowing for continuously stable neural activity. Using computational modelling we demonstrate that early in development excitatory short-term depression quickly stabilises neural activity, even in the face of strong, unbalanced excitation. We introduce a model of the commonly observed developmental shift from depression to facilitation and show that neural activity remains stable throughout development, while inhibitory synaptic plasticity slowly balances excitation, consistent with experimental observations. Our model predicts changes in the input responses from phasic to phasic-and-tonic and more precise spike timings. We also observe a gradual emergence of synaptic working memory mediated by short-term facilitation. We conclude that the developmental depression-to-facilitation shift may control excitation-inhibition balance throughout development with important functional consequences.

## Introduction

Short-term synaptic plasticity is a hallmark of synaptic function. It refers to transient and fast changes in synaptic efficacy in the range of a few milliseconds up to several seconds ^1–3^. Different short-term plasticity (STP) profiles regarding the direction and time scale of change are found across cell types ^4–7^, brain regions ^8–12^, and throughout development ^8–10,13–15^. For example, excitatory synapses from pyramidal cells in cortex are predominately short-term depressing in young animals, whereas adult synapses exhibit short-term facilitation ^8^. Conversely, inhibitory synapses from cortical fast-spiking inhibitory interneurons are short-term depressing throughout development ^4,6,7^. Functionally, STP is known to homeostatically control synaptic transmission and firing rates in neuronal networks on millisecond timescales ^16–18^. However, it has remained unclear what is the combined impact of long-term and short-term plasticity for homeostatic control in neural circuits.

Recent studies suggest that long-term inhibitory plasticity (ISP) ^19–23^, acting on the time scale of minutes to hours, is also responsible for homeostasis, by way of establishing and maintaining excitation-inhibition balance, limiting the destabilising effects of its excitatory counterpart ^24,25^. However, the stabilising effects of co-tuning excitatory and inhibitory tuning curves, the hallmark of inhibitory synaptic plasticity, can only be observed in adult animals. In young animals, a tight excitation-inhibition balance has not yet formed and receptive are often unbalanced ^25,26^. Despite this lack of detailed excitation-inhibition tuning, experimental observations consistently show that neural circuits exhibit stable firing activity at all stages of development ^27–30^. Here, we hypothesise that short-term plasticity provides the homeostatic control needed in young animals for healthy neural activity.

Using computational models, we show how short-term plasticity can complement and even control the expression of inhibitory long-term plasticity, thus acting as a gating mechanism for the emergence of excitation-inhibition balance across development. In particular, we show that short-term depression is critical to maintain stable neural activity even with flat inhibitory tuning curves in young animals ^25^. Further, the gradual shift to short-term facilitation, as observed throughout development ^8–10,13–15^ allows for excitatory-inhibitory balance to emerge. We show that this developmental control of STP shapes the properties of neuronal dynamics, making neural responses more diverse and postsynaptic spike timings more precise over the course of maturation. Finally, the maturation of STP in our model leads to synapse-based working memory properties in a EI balanced neuron model.

## Results

Changes in short-term plasticity (STP) are a hallmark of neural development ^8,12,31^, but their impact on neuronal dynamics has remained unclear. Here, we study the effects of short-term plasticity in congruence with long-term inhibitory plasticity in a developing neuron model, and show that STP can play a crucial role in young neurons, compensating for a lack of inhibitory tuning. Moreover, gradual change of excitatory STP from depression to facilitation over development allows for excitatory-inhibitory balance to develop in the neuron while guaranteeing stable response properties.

To investigate these effects, we built a model of a simple feedforward network with a single conductance-based integrate-and-fire neuron receiving inputs from 800 excitatory and 200 inhibitory afferents ^21^. To emulate heterogeneous inputs we modelled eight different pathways (Fig. 1a) each with 100 excitatory and 25 inhibitory synapses, whose activity is determined by a time-varying rate signal (Methods). Excitatory and inhibitory synapses were modulated by short-term plasticity, consistent with experimentally observed profiles in young and adult mice ^8–10,12–14,31–35^. Inhibitory synapses additionally experienced long-term plasticity (ISP) ^19,22^. Excitatory afferents were tuned according to experimentally observed receptive fields, while inhibitory baseline weights were initially flat (Fig. 1b, see also ^25^)

**Figure 1.**
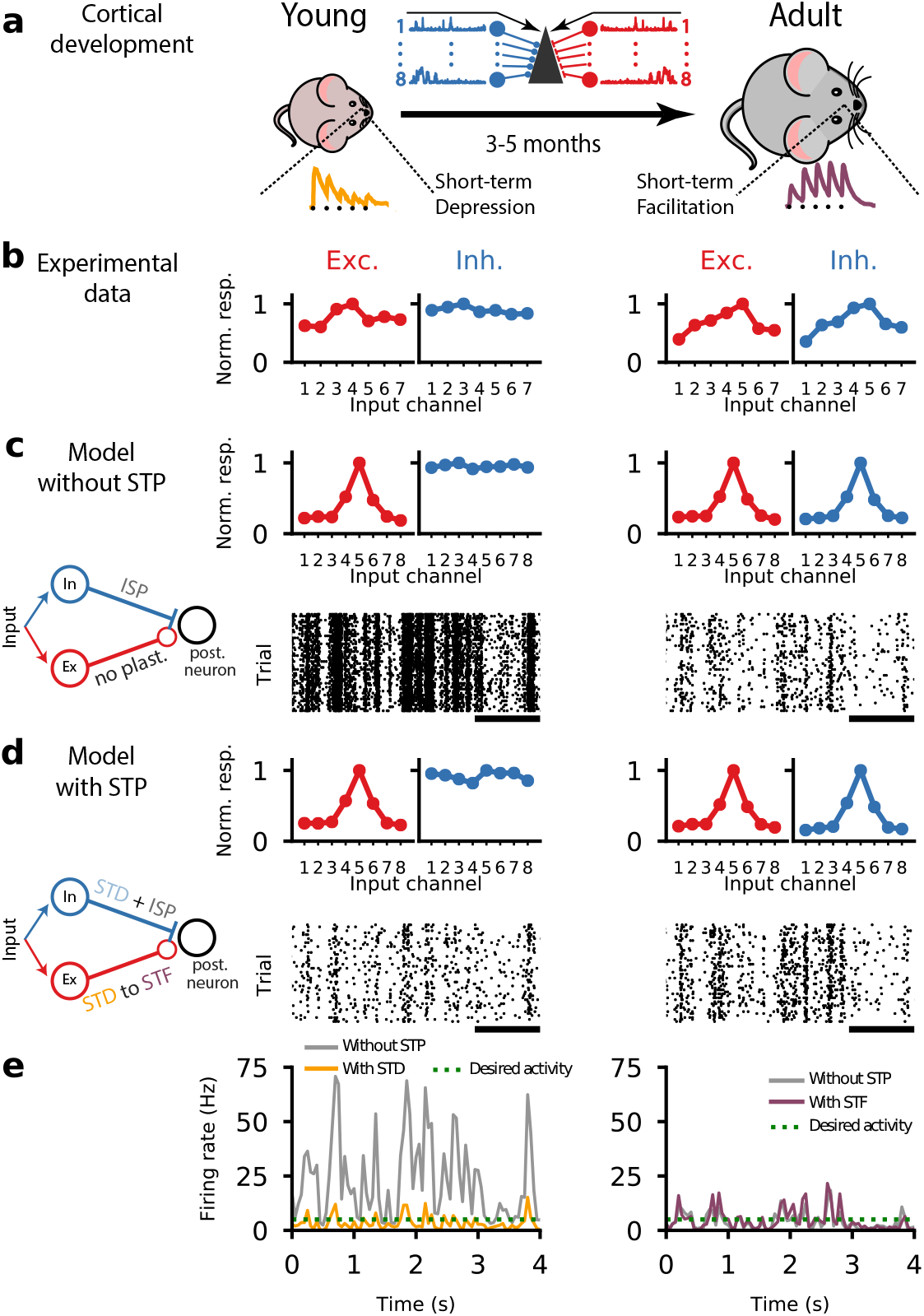
A cortical circuit with short-term synaptic plasticity exhibits healthy neural dynamics in both young and adult conditions. **(a)** Schematic of animal development from young with short-term depression (left) to adult with short-term facilitation (right) at excitatory synapses as observed experimentally ^8–10,13,14^. Traces of short-term synaptic plasticity (STP) for depression (orange) and facilitation (purple) ^8^. In the middle is a schematic of the feedforward neural circuit with eight independent input channels, each with an excitatory (red) and an inhibitory (blue) group synapsing onto a postsynaptic neuron (Fig. S1). **(b)** Inhibitory tuning does not mirror excitatory tuning in young animals (left). Once animals reach adulthood, a precise excitation-inhibition (EI) balance can be observed. Panels adapted from Dorrn et al. ^25^. **(c)** Computational model with long-term synaptic plasticity in inhibitory synapses (ISP; see inset) started from unbalanced excitation-inhibition (top left) and gradually developed EI balance (top right). Neuron with unbalanced excitation-inhibition showed high activity (~20 Hz; bottom left), which was gradually reduced through ISP (~4.5 Hz; bottom right). Bottom raster plots represents postsynaptic spiking activity. **(d)** A computational model with both ISP and STP started from unbalanced excitation-inhibition (top left) and gradually developed EI balance (top right). Neuron with unbalanced excitation-inhibition shows low/healthy firing activity (~4.5Hz; bottom left) throughout development (~4.5Hz; bottom right). Bottom raster plots represents postsynaptic spiking activity. **(e)** Firing rates of a model without STP (solid gray line) and a model with both ISP and STP in young (left, solid orange line) and adult (right, solid purple line) conditions. Desired activity (dashed green line) represents baseline firing rate as observed experimentally ^27–30^.

Inhibitory long-term synaptic plasticity working on a time-scale of hours has been suggested to underlie excitation-inhibition (E-I) balance in cortical networks ^19,21,22^. The slow nature of long-term synaptic plasticity is consistent with the gradual and slow development of E-I balance over multiple days from young to adult animals ^25^ (Fig. 1b). However, the lack of detailed balance in young animals could lead to unstable, unnaturally high activity (Fig. 1c,e). Increased learning rates, on the other hand, lead to unstable learning ^36,37^.

Short-term plasticity can offer an elegant solution to maintain low firing rates throughout development. To this end, we added experimentally observed ^4,6,7^ short-term depression to all afferent synapses using a standard Tsodyks-Markram model ^16^ (Methods). In contrast with the ISP-alone model, the addition of an appropriate STP profile that features short-term depression at the excitatory synapses, led to lower firing rates in the ‘young’ model, despite unbalanced excitation-inhibition (Fig. 1d,e).

Notably, the low postsynaptic firing rates that resulted from short-term depression in the excitatory afferents effectively prevented long-term plasticity from tuning inhibitory tuning curves as has been observed in adult animals (Fig. 1b; ^25^). As we will see below, the shift of short-term plasticity profiles over the course of development ^8,12,31^ allowed the gradual tuning of inhibition in ageing animals.

### Gradual depression-to-facilitation shift enables stable activity over development

Next we studied how the developmental changes of short-term depression (STD) to short-term facilitation (STF) in excitatory synapses ^8–10,12–14,31–35^ may aid the tuning of inhibitory synapses by way of long-term plasticity, and provide stable postsynaptic firing rates throughout the process.

To simulate ageing in our model, we devised an algorithm that slowly changed the STP parameters between young and adult profiles fitted to experimental data (Fig. 2a; Methods). The algorithm monitored average postsynaptic firing over sliding windows of 500 ms. When rates were stable and low, excitatory STP parameters were modified by a small amount towards facilitation (see Methods and Figs. S2–S4 for variations). For computational reasons we used a total simulation time of 8 hours to model development, but the exact temporal frame does not qualitative change our results (data not shown).

**Figure 2.**
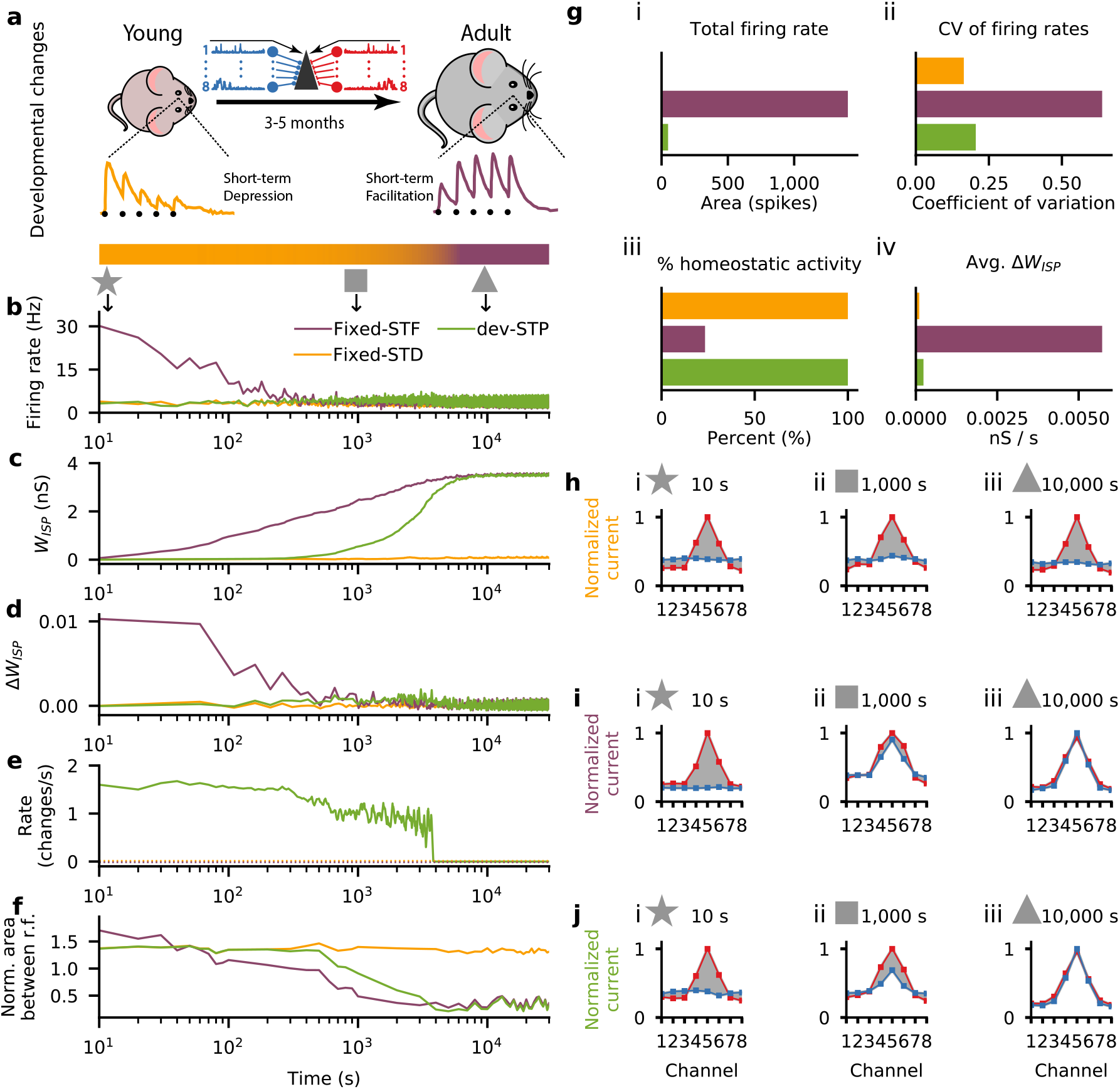
Gradual short-term plasticity shift maintained stable firing rates while detailed EI balance developed. (**a**) Schematic of our developmental short-term plasticity (STP) model (cf. Fig. S1); top: young and adult STP (as in Fig. 1); bottom: gradual changes in STP from depressing to facilitating dynamics (orange and purple respectively, in log-scale as in b-f). (**b**-**f**) Different variables of the model across simulated development for three different models: fixed short-term depression (fixed-STD, orange), fixed short-term facilitation (fixed-STF, purple) and developmental model with gradual changes in STP (dev-STP, green line). Note x-axis on log-scale. (**b**) Receiver neuron firing rate. (**c**) Mean inhibitory weight. (**d**) Mean changes in the weight of the inhibitory synaptic afferents. (**e**) Rate of STP change (note that both fixed-STF and STD remain fixed, shown as dashed lines). (**f**) Area between normalised excitatory and inhibitory tuning curves (cf. h-j) during the course of simulated development. A normalised area close to 0 represents a perfectly balanced neuron. (**g**) Additional statistics for the three models. (i) Total neuronal activity calculated using the area between the firing rate in (b) and the desired target rate of 5 Hz. (ii) Average coefficient of variation of the firing rates across simulated development (cf. (b)). (iii) Percent of time spent under homeostasis (i.e. at the desired firing rate; cf. (b)). (iv) Average change in inhibitory weights (cf. (d)). (**h**-**j**) Snapshots of excitatory and inhibitory tuning curves across three points in simulated development: 10s (star), 1000s (square) and 10 000s (triangle). Shaded gray area represents difference between excitatory and inhibitory tuning curves (cf. (f)). (**h**-**j**) Excitatory (red) and inhibitory (blue) postsynaptic tuning curve for the fixed-STD (h), fixed-STF (i) and dev-STP models (j).

The developmental STP model (dev-STP) maintained a healthy level of firing activity throughout the simulation (i.e. approximately 5Hz) while a tight excitation-inhibition balance in the circuit developed (Fig. 2b). As controls, we considered two other models in which STP was fixed either at STD (fixed-STF) or STF (fixed-STF). The fixed-STF scenario exhibited high and more variable firing rates before ISP was able to balance the postsynaptic neuron and lower the firing rates (Fig. 2b,g; Fig. S5). On the other hand, the fixed-STD scenario was able to maintain homeostatic balance throughout the simulation (Fig. 2b,g), but did not develop a tightly balanced inhibitory receptive field (Fig. 2f,h).

Although the developmental STP and fixed-STF models converged to the same mean inhibitory weights (Fig. 2c), the fixed-STF scenario led to substantially higher firing rate variability during development, and large, somewhat erratic weight changes (Fig. 2g,d). In contrast dev-STP maintained relatively small weight changes throughout development (Fig. 2d). Finally, while the initial changes of receptive field in the fixed-STF scenario arose quickly, the time of convergence was similar to the dev-STP model (Fig. 2f,i,j), because long-term inhibitory plasticity in the dev-STP scenario sped up dramatically as facilitation developed (Fig. 2b-f). In the dev-STP model, ISP evolved the inhibitory tuning to match excitation stepwise (Fig. 2f), incrementally handing over control of the target firing rate to inhibition, which ensured postsynaptic activity remained relatively low (Fig. 2b). This means that each increase in the excitatory efficacy through strengthened STF was matched by an increase in the inhibitory efficacy through ISP, until inhibition was fully tuned and the excitatory synapses reach their adult profile of short-term facilitation.

The dev-STP model was able to maintain the neuron in a (globally) balanced state throughout development while allowing inhibition to gradually mirror the excitatory tuning. In line with experimental *in vivo* observations in rat auditory cortex across development ^25^ inhibitory tuning curves were initially flat (Fig. 3a). In the adult neuron, both model and experiment showed E-I balance. Using the same linear correlation analysis as in the experimental work, we confirmed that excitatory and inhibitory responses in ‘young’ models were not correlated, but became strongly correlated in the adult profile (Fig. 3b).

**Figure 3.**
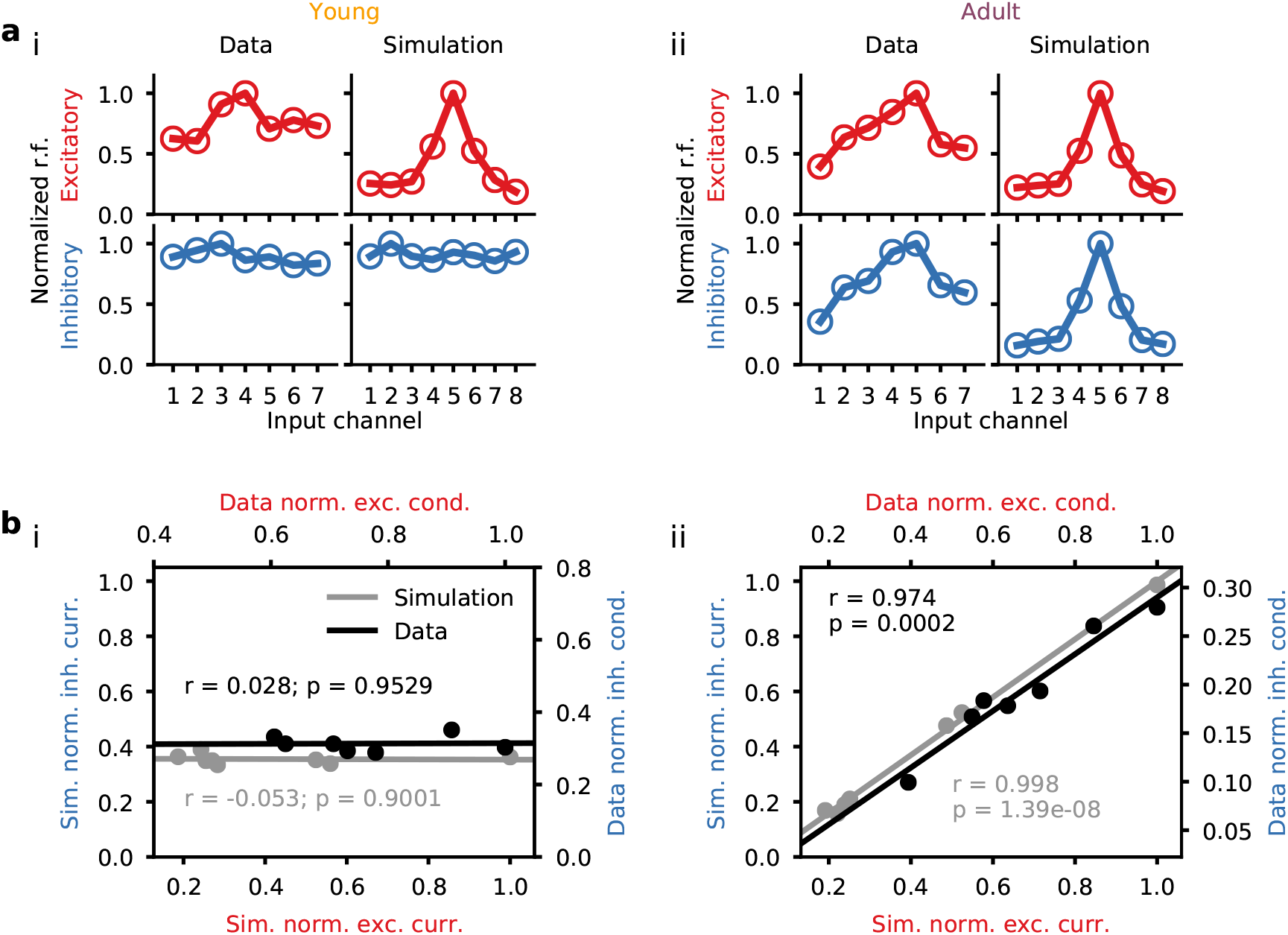
Depression-facilitation shift captured inhibitory receptive field development. (**a**) Comparison of experimentally observed and simulated (dev-STP model) excitatory and inhibitory tuning curves, for both young (i) and adult (ii) conditions. (**b**) Excitatory-inhibitory responses for model (gray) and experiments (black). Different dots represent different tone frequencies in the data and different input channels in the model. Lines represent linear correlation between excitatory and inhibitory responses in both model (gray) and experiments (black). Experimental data reproduced from Dorrn et al. ^25^.

### Developmental changes in STP shape signal dynamics and transmission

In line with the establishment of detailed balance ^21^, the postsynaptic firing rates in the dev-STP model were initially more correlated with the fixed-STD model, and gradually became more correlated with the fixed-STF model (Fig. 4a,b,c; Fig. S5). Across all input channels we found a gradual decrease of input-output correlation (Fig. 4d)). This was largely due to the fact that the output responses became less correlated with the preferred channel versus the non-preferred channels (Fig. 4e).

**Figure 4.**
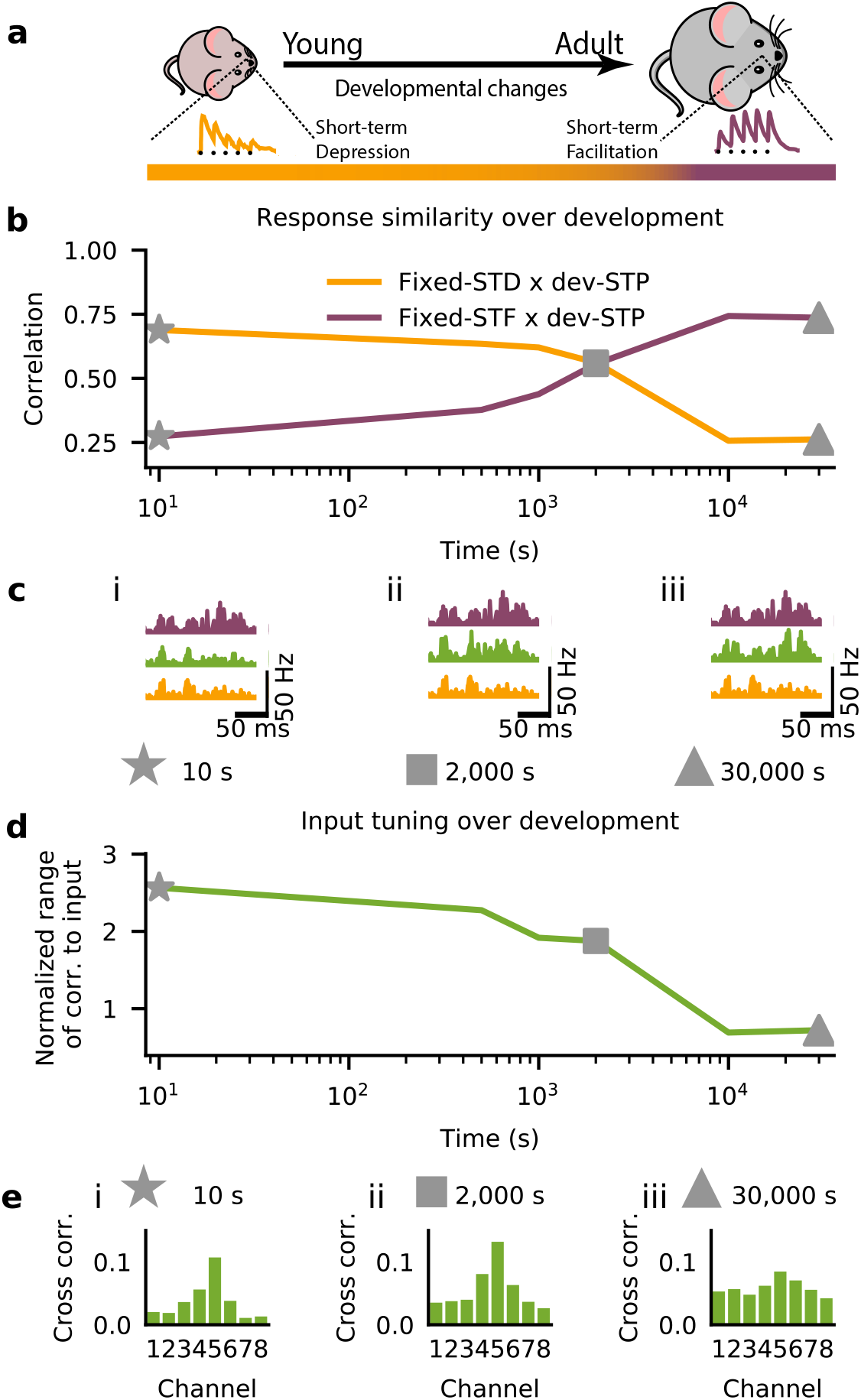
Input-output response correlations over development. (**a**) Schematic of the modelled development from young with depressing synapses (left) to adult facilitating synapses (right). Bottom color bar indicates the gradual shift in STP (as in Fig. 2). (**b**) Correlation of the dev-STP model response profiles to that of the fixed-STD (orange) and fixed-STF (purple) scenarios during development. (**c**) Example output responses (cf. Fig. S5) for the fixed-STD (orange), fixed-STF (purple), and dev-STP (green) models at three points in simulated development (i: 10s, stars; ii: 2000s, squares; iii: 30000s, triangles). (**d**) Normalised range of correlation to input (Methods). (**e**) Example of output correlations at specific times during the course simulated development (same timings as in c). Results shown here were averaged over 50 trials.

Another functional consequence of the changes in short-term dynamics could be observed in the phasic and tonic stimulus responses profiles. Transient (phasic) and steady state (tonic) neural activity has been observed in sensory cortical circuits as part of their stimulus response repertoire ^27,33,38,39^. We examined these properties by probing the neuron responses using a step input stimulus (see Methods) (Fig. 5b) to the preferred input channel (channel 5), simulating the sudden presence of a strong sensory feature. We defined the phasic response as the average activity over the first 50ms after stimulus onset, and the tonic response as the average rate over the remaining stimulus duration (200ms). Over development, the average phasic activity of the circuit decreased, while the tonic activity increased (Fig. 5b; Fig. S6). These changes in the dynamics are a direct consequence of the gradual change from depressing to facilitating synapses, interacting with the strengthening inhibition. The shift in tonic and phasic responses to a single stimulus also affected subsequent input responses when using two paired step inputs (Fig. 5d inset; Methods). This interaction between subsequent responses was largest for the phasic response, which grew substantially over development, as seen by the increasing ratio of firing rate between the two stimuli (Fig. 5d,e). On the other hand, the tonic response decreased, but only slightly.

**Figure 5.**
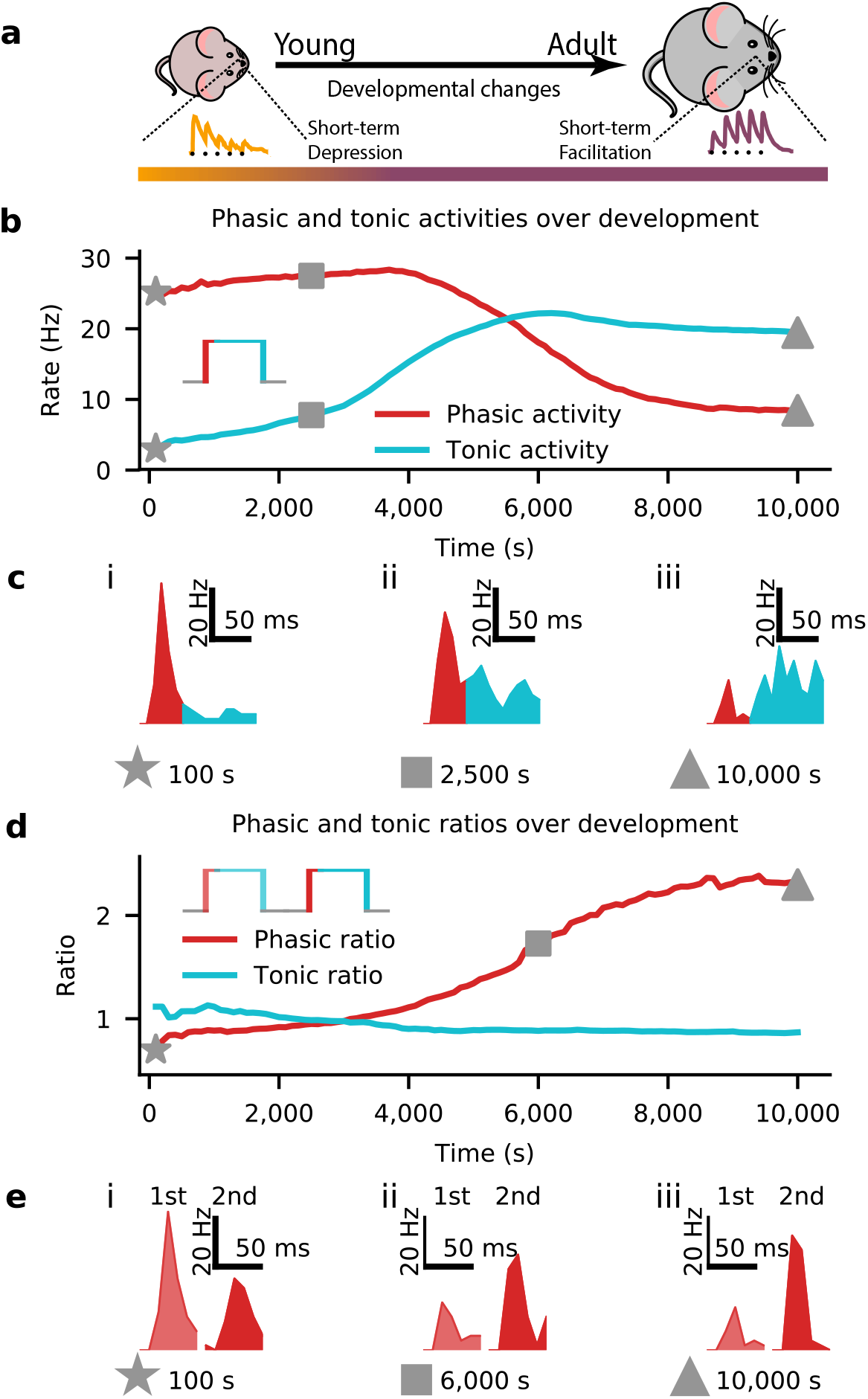
Developmental STP shaped tonic and phasic input-output transmission. (**a**) Schematic of the modelled development from young with depressing synapses (left) to adult facilitating synapses (right), as in previous figures. (**b**) Average phasic (red) and tonic (blue) postsynaptic firing rates for a step-input of 150Hz (inset; cf. Figs. S5,S6). (**c**) Example output responses for the phasic (red) and tonic (blue) activities at three points during development. (**d**) Ratios of the average phasic (red) and tonic (blue) firing rates between two consecutive step stimuli (inset). (**e**) Examples of responses to the first (light red) and second (dark red) phasic activities in response to the double step input stimulus at specific points during development. Results shown here were averaged over 50 trials.

We also investigated the phasic response to a step stimulus on very short time scales (Fig. 6a), specifically focusing on the temporal jitter of the first evoked spike (Fig. 6b). In line with previous experimental observations of reduced jitter over development ^25^, we observed substantially more stimulus-locked spike times in the adult model than in the young model (Fig. 6c,d). The young scenario showed higher normalized jitter (Methods) than the adult scenario across all stimulus strength, and particularly when the background activity before stimulus onset was low (Fig. 6e).

**Figure 6.**
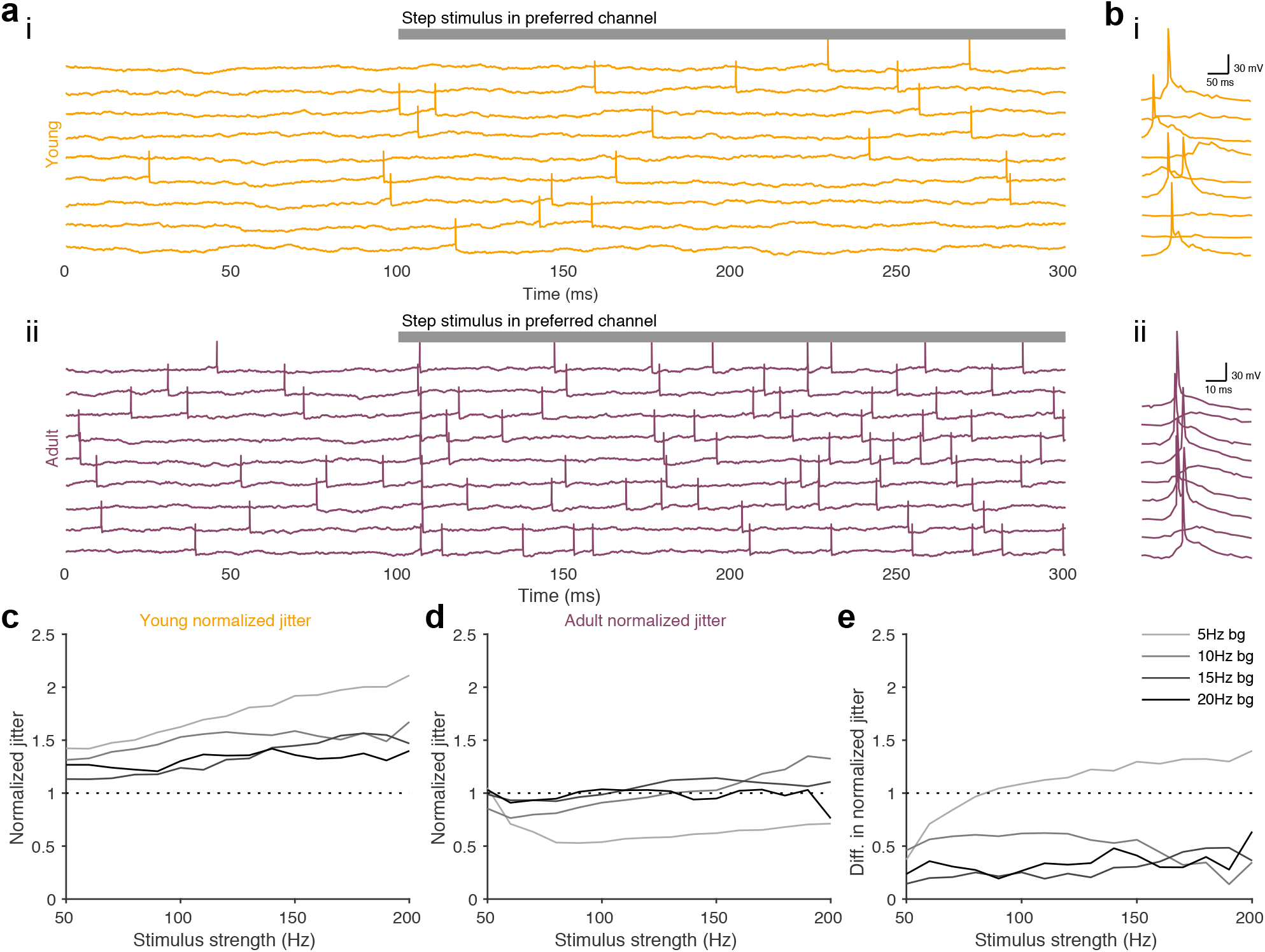
Adult STP improves temporal precision of postsynaptic spikes. (**a**) Examples of postsynaptic voltage responses with preferred-channel input for both young STP model (i) and adult STP model (ii); gray bar at top represents time during which preferred channel is stimulated. (**b**) Stimulus evoked responses in *in vivo* recordings across a few trials in young (i) and adult (ii) animals. Panels adapted from Dorrn et al. ^25^. In (a,b) the background firing rate is 5 Hz. (**c,d**) Normalized jitter of postsynaptic spikes in the young (c) and adult (d) model for different background firing rates (denoted by different shades of gray; see Methods). (**e**) Difference between normalized jitter of young STP model (c) and adult STP model (d).

### Developmental STP enables working memory properties in a balanced neuron

Finally, we also investigated the longer term effects of changing STP over development with regard to its implications for short-term memory. Short-term plasticity has recently been proposed as a substrate for working memory ^40,41^, owing to the fact that STF can promote increased response to previously displayed stimuli. Here, we tested these ideas in the dev-STP model, by comparing the responses to “recall” stimuli that were or were not preceded by a “preloaded” stimulus.

Models with no STP mechanism, as well as the ‘young’ dev-STP model showed identical firing rates during the recall period (Fig. 7a,b) independently of whether they had experienced a preloaded stimulus or not. In other words, the ‘young’ model could not rely on silent working memory traces. The ‘adult’ dev-STP model, on the other hand, showed substantially higher firing rates during the recall period (Fig. 7c,d) when the recall stimulus was preceded by a preloaded cue that activated the short-term facilitation in excitatory synapses. Dev-STP thus allowed the neuron to gradually utilise this silent working memory mechanism in a neuron with EI balance (Fig. 3a,b, Fig. 7e,f).

**Figure 7.**
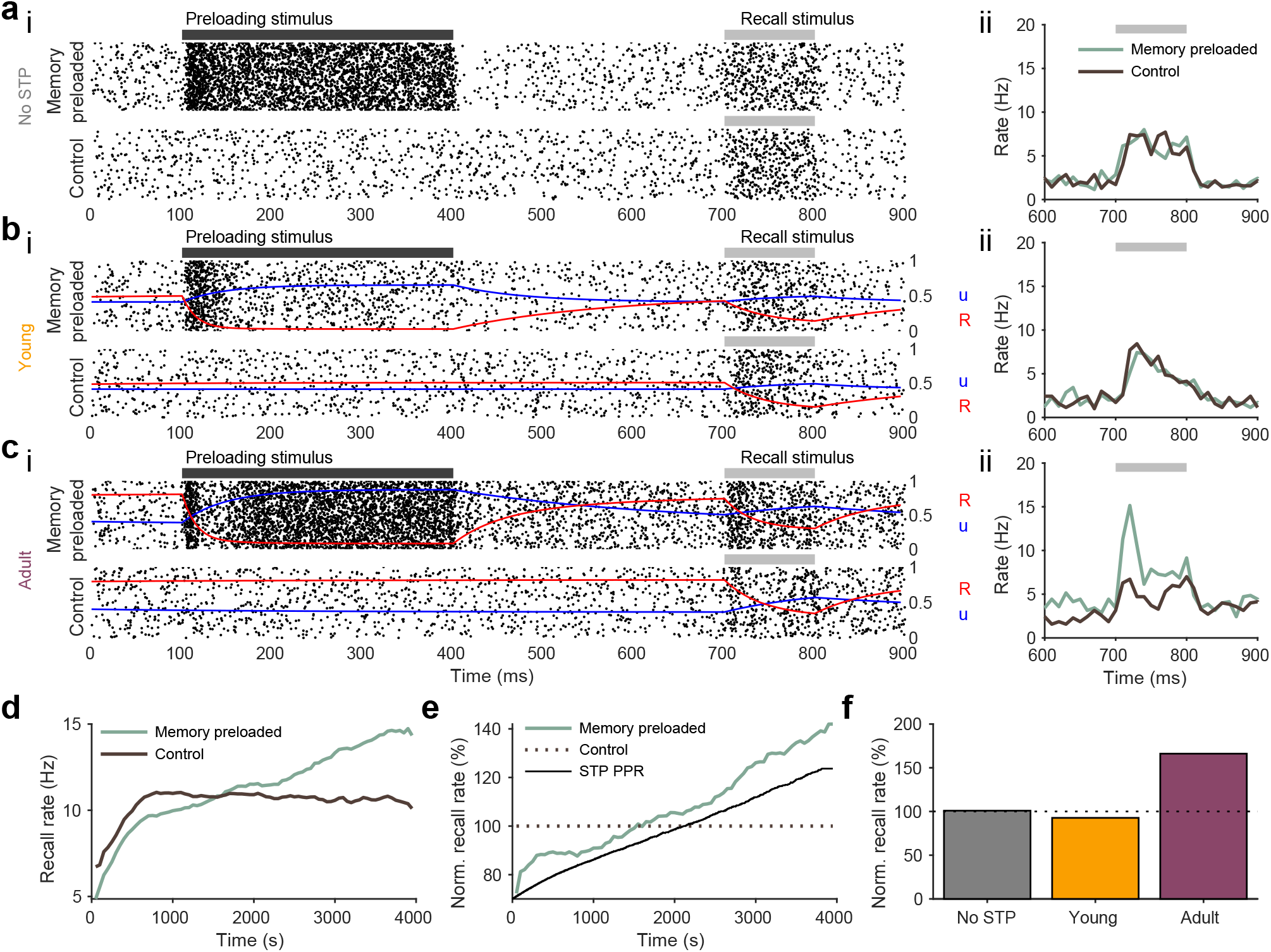
Gradual emergence of synaptic-based working memory over development. (**a-c**) Raster plot of a working memory test (WMT, i-top)) with a preloaded stimulus and subsequent recall stimulus (black and gray bars respectively) compared with rasterplot of trials without the preloaded stimulus (i-bottom). Average firing rates (ii) for both memory preloaded (light green) and control conditions (dark brown). (**a**) WMT in a model with only inhibitory synaptic plasticity (i.e. no STP) (**b**) WMT in a model with young STP profile. (**c**) WMT in a model with adult STP profile. (**d**) Firing rates during the recall period with (light green) or without (dark brown) preloaded stimulus. WMTs were preformed every 50 seconds during dev-STP development simulation (cf. Fig. 2) as STP changes from depression to facilitation at excitatory synapses. We only highlight the first 4000s of the simulation as after this point STP become minimal. (**e**) Normalised recall firing rates to the average firing rate of the control case (i.e. without memory preloading). The STP paired-pulse ratio (black) measuring the STP strength of the excitatory synapses for this period is also plotted as reference. (f) Normalized recall rate for three model conditions: no STP (gray), young STP (orange), and adult STP (purple).

## Discussion

It has been widely observed that short-term synaptic dynamics of the cortex change from depressing to facilitating throughout the course of development ^8–10,12–14,31–35^. Here, we show that this commonly observed shift in STP may interact with long-term plasticity at inhibitory synapses to form the fundamental architecture of neuronal processing. According to our model, short-term depressing synapses could help to stabilize neural networks in the absence of properly tuned inhibition in young animals (Fig. S5). A gradual change from short-term depression to facilitation then allows for stable dynamics throughout development while inhibitory synaptic plasticity-mediated, detailed excitation-inhibition balance can emerge (Fig. 2). In addition to this stabilising interplay, we show that the developmental maturation of STP also shapes signal processing, by allowing for more temporally precise coding (Fig. 6), and the emergence of synaptic working memory (Fig. 7).

There are currently two dominant views on how changes in STP throughout development may arise. One view is that these changes are caused by sensory experience ^32^; the other view poses that these are hard-wired, pre-programmed changes ^13^. Our developmental STP model suggests a way to reconcile these two views, in that both the sensory-dependent ^32^ and non-sensory-dependent ^13^ changes observed experimentally may be simply caused by changes in the neural baseline activity. However, although we have modelled changes in STP as a function of neural activity, it is in principle possible to allow for these changes to be purely hard-wired and continuous (cf. Fig. S4). In our hands, the latter mode, i.e. unilateral maturation of STP without heeding the co-development of inhibitory tuning curves, can also lead to stable development (Fig. S4), but this requires fine tuning of a STP change interval, and additional experimental work remains to be done to further study this scenario.

Our work highlights how developmental-STP may shape temporal aspects of synaptic transmission. In particular, our model predicts that young animals primarily encode stimuli with transient, phasic activity, whereas adult animals may transmit both phasic transients and sustained tonic rates equally well. Interestingly, both modes of transmission have been observed in sensory cortices ^27^ at different developmental stages. In our model we have assumed that STP changes at all excitatory synapses happen in lockstep over development. However, in the brain not all synapses are modified coincidentally ^8–10^, and it is possible that this degree of variability gives a tighter homeostatic control throughout development.

We have focused on long-term inhibitory synaptic plasticity, but excitatory synapses also undergo long-term synaptic plasticity. Importantly, long-term excitatory synaptic plasticity also changes the short-term synaptic dynamics ^18,42–44^. It is possible that the gradual changes of STP at excitatory synapses that we have considered here are mediated by long-term excitatory plasticity. In future work it would be interesting to explore the effects of long-term excitatory plasticity with realistic inputs in conjunction with inhibitory synaptic plasticity as a potential model for developmental STP ^19,36,45^.

Our model shows a gradual increase in temporal precision of spiking over development, consistent with experimental observations in the auditory cortex of rats ^25^, suggesting that STP maturation plays an important role in temporal encoding ^46–50^. Our findings add to the growing experimental literature showing that inhibition-excitation balance sharpens spike timings ^25,49,51,52^.

Working memory is traditionally thought of as being a property of recurrent neural network dynamics in the prefrontal cortex. However, forms of working memory are also known to exist in sensory cortices ^53,54^. Moreover, short-term facilitation has been proposed as a biologically plausible mechanism of working memory at the synaptic level ^40,55^. We have shown that working memory-like properties in line with previous theoretical work ^40^ gradually emerge in our model as short-term facilitation becomes more dominant. Moreover, we show here that retaining EI balance does not interfere with this type of silent working memory. Our results suggest that silent synaptic working memory properties are more likely prevalent in adult cortex, potentially enabling animals to retain information about the recent past even in sensory cortices.

Finally, dysfunctions in the regulation of excitation-inhibition balance underlie numerous neurological disorders ^56–65^. In our model we show that short-term plasticity can dynamically control the expression of long-term inhibitory synaptic plasticity, thus modulating E-I balance. Maldaptive developmental STP should thus be reflected in E-I malfunction. Interestingly, this is supported by disease animal models, in which STP and excitation-inhibition balance are both altered in animal models of dysplasia ^66,67^.

Overall, our results suggest important functional roles for the commonly observed shift in STP during development.

## Acknowledgements

We would like to thank the Vogels Lab for feedback on an earlier version of this manuscript. D.W.J. was supported by a Marshall Scholarship and a Clarendon Scholarship. R.P.C. and T.P.V. were supported by a Wellcome Trust and Royal Society Sir Henry Dale Fellowship (WT 100000), a Wellcome Trust Senior Research Fellowship (214316/Z/18/Z), and an ERC Consolidator Grant (SYNAPSEEK).

## Supplementary material

### Materials and Methods

#### Neuron models

In this study, we used a conductance-based integrate-and-fire neuron model for simulations ^68^. In this model, the membrane voltages are calculated following

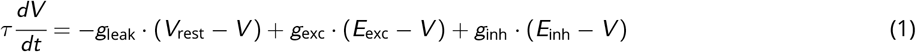

where *V* is the membrane potential of the neuron as a function of time *t*, *τ* is the membrane time constant, *V*_rest_ is the resting membrane potential, *E*_exc_ is the excitatory reversal potential, and *E*_inh_ is the inhibitory reversal potential. Our neuron parameters are the same as in Vogels and Abbott ^68^. In particular, we used a membrane capacitance, *C*, of 200pF with membrane resistance, *R*, of 100MΩ, which gives a membrane time constant *τ* = 20ms. *g*_exc_ and *g*_inh_, expressed in the units of the resting membrane conductance, are the synaptic conductances, and *g_l_* is the leaky conductance. The synaptic conductances are modelled as 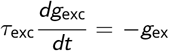 and 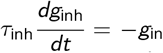where *τ*_exc_ and *τ*_inh_ are the synaptic time constants for the excitatory and the inhibitory conductances, respectively. When the neuron receives a presynaptic action potential, its conductance increases by *g*_exc_ → *g*_exc_ +*w*_exc_ or *g*_inh_ → *g*_inh_+*w*_inh_ for excitatory and inhibitory synapses, respectively. The model parameters used are summarized in Table 1.

**Table 1.** Parameter values for conductance-based leaky integrate-and-fire model.

| Parameter | Value |
| --- | --- |
| $\tau$ | 20.0ms |
| $R$ | 100.0MΩ |
| $C$ | 200.0pF |
| $g_{\text{leak}}$ | 10.0nS |
| $\tau_{\text{exc}}$ | 5.0ms |
| $\tau_{\text{inh}}$ | 10.0ms |
| $E_{\text{exc}}$ | 0mV |
| $E_{\text{inh}}$ | -70mV |
| $V_{\text{rest}}$ | -60mV |
| $V_{\text{thresh}}$ | -50mV |
| $V_{\text{reset}}$ | -60mV |
| $\tau_{\text{refrac}}$ | 4ms |
| $w_{\text{exc}}$ | 0.5ns |
| $w_{\text{inh}}$ | 0.1ns |

### Synaptic plasticity models

We used both short-term plasticity and long-term inhibitory synaptic plasticity models in our work. Both were calculated separately in the simulations and combined as explained below.

#### Short-term synaptic plasticity (STP)

Short-term plasticity was used in the simulations following the model defined by ^5,69,70^ following

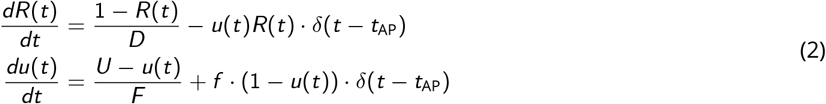

where *R* models vesicle depletion and *u* models the presynaptic release probability. Every presynaptic spike at *t*_AP_ causes a decrease in *R*, the number of vesicles available by *uR*, which then recovers exponentially to its baseline value of 1 with a time constant *D*. At the same time every presynaptic spike at *t*_AP_ also causes an increase in the release probability *u* by *f* · (1 − *u*(*t*)) (where *f* is the facilitation rate) and recovers exponentially to its baseline *U* with a time constant *F*. Finally, the postsynaptic potential, or the weight of the STP component for a synapse exhibiting STP at time *t* is computed as *w*_STP_(*t*) = *AR*(*t*)*u*(*t*), where *A* is baseline amplitude factor. In simulations, the initial value of *u* is set to *U*, and the initial value of *R* is set to 1. We used the four-parameter version of the TM model (*D*, *F*, *U*, *f*) as it provides an overall better fit of short-term dynamics data ^70^.

#### STP model fitting

We found STP parameters which produced excitatory STP paired-pulse responses (PPRs) that matched those found in experiments for young and adult animals. Specifically, we used the STP PPRs observed by Reyes and Sakmann ^8^, with excitatory STP PPRs of 0.7 and 1.24 for young and adult animals respectively. In order to find STP parameter values that matched these PPRs, we interpolated between strong STD and strong STF parameter values ^70^ (Fig. S1e). Using this interpolation we then calculated the PPR across all parameter sets. We use these PPRs to compared with experimental data from young and adult animals as observed in Reyes and Sakmann ^8^. Finally we used least squares to obtain STP parameters that best matched the data in both young (STD) and adult conditions (STF) (see Table 2).

**Table 2.** STP parameter values. Paired-pulse ratio (PPR) is given by dividing the second postsynaptic response by the first.

| Synaptic dynamics | $D$ (s) | $F$ (s) | $U$ | $f$ | PPR |
| --- | --- | --- | --- | --- | --- |
| Depression | 0.3134 | 0.0798 | 0.3917 | 0.062 | 0.70 |
| Facilitation | 0.0845 | 0.2959 | 0.1973 | 0.1168 | 1.24 |

#### Inhibitory synaptic plasticity

Long-term inhibitory synaptic plasticity (ISP) is implemented in all inhibitory synapses in all simulations unless otherwise specified. We used the same model as Vogels et al. ^21^. In this model, each synapse *i* has a presynaptic trace *x_i_*, which increases with each spike by *x_i_* → *x_i_* + 1 and decays exponentially following 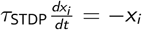. Then, the synaptic weight of a given synapse following pre- or postsynaptic spikes are updated by

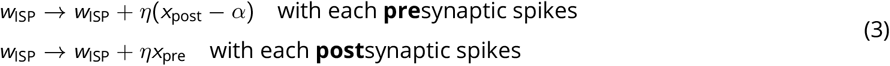

where *η* is the learning rate, *α* = 2 · *r*_target_ · *τ*_STDP_ is the depression factor, where *τ*_STDP_ = 20ms is the STDP time constant, and *r*_target_ = 5Hz is a constant parameter that defines the target postsynaptic firing rate. In simulations, the initial values of *w*_ISP_ is set to zero.

#### ISP with STP

In our simulations, ISP is combined with STP in some cases at the inhibitory synapses. In these cases, the total synaptic weight *w*_inh_ is computed as the product of the STP and ISP weight components at the time of the postsynaptic spike 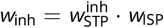 while the excitatory weight was given by 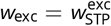.

### Simulations

#### Input signals and connectivity

To model the neural responses with naturalistic inputs we used 8 independently generated traces of low-pass filtered, half-wave rectified white noise signals. Each of the 8 independent channels represents a signal pathway, and consists of 100 excitatory neurons and 25 inhibitory neurons, giving a total of 1000 presynaptic neurons ^21^. All presynaptic neurons synapse onto a single postsynaptic neuron with a total of 1000 synapses, 800 excitatory and 200 inhibitory.

As in Vogels et al. ^21^ for each of the 8 channels, we generated its time-varying rates iteratively as 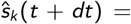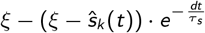 where 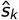 is the *k*-th signal, *ξ* ∈ [−0.5, 0.5] is drawn from a uniform distribution, *dt* = 0.1ms is the simulation time step, and the filtering time constant is *τ_s_* = 50ms. We normalized all rates to a preferred firing rate of 100Hz, and negative values were remove and replaced with a background activity level of 5Hz.

These traces represent the firing rates across time of each of the 8 input signal channels (see examples in Fig. S1b). We used these rates as seeds to generate Poisson spike trains for each of the eight channels. These inputs were used in the simulations shown in Figures 1, 2, and S5.

#### Developmental and fixed STP

When simulating dev-STP, we first found the STP parameters whose paired-pulse ratio (PPR, i.e. EPSP_2_/EPSP_1_) best matched experimental data ^8^. To this end, we started with STP parameters which give strong depression and strong facilitation ^70^. Next, we conducted a parameter sweep of the STP parameters from strong depression to strong facilitation using a dense linear space between these two conditions. We then simulated 50 Poisson input spike trains at 35Hz ^8^, calculated the average PPRs of each train for all STP parameters. We then used the STP parameter values that best matched those of Reyes and Sakmann ^8^ for our simulations. These parameter values are summarized in Table 2.

#### Calibrating the parameters for dev-STP

Using the STD and STF parameters given in Table 2, we then calculated a set of 3600 parameter values spaced logarithmically between the STD and the STF parameter values. Log interpolation was used instead of linear interpolation because a marginal change towards facilitation generates a higher marginal change in PPR when closer to facilitation than to depression. For each of the 3600 STP parameter values, each time we changed STP parameters, we normalized the STP magnitude parameter *A* to equal

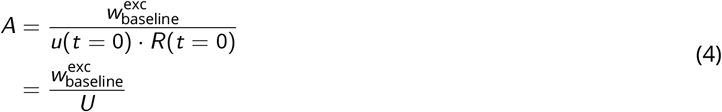

where 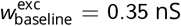 is the baseline excitatory weight. This normalization fixed the amplitude of the first PSP to the same value, regardless of the STP parameters, thus keeping the baseline weight of excitatory synapses the same throughout development during the simulation (see below for alternative normalizations). Note that the initial value of *u* is set to *U*, the initial value of *R* is set to 1, and the total excitatory weight for a first pre-synaptic spike is given by

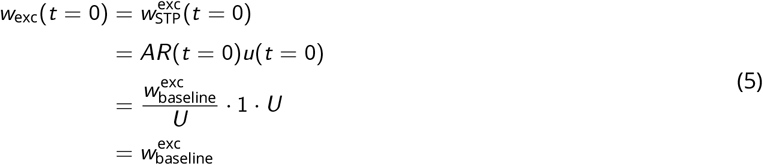

regardless of the STP parameters, thus the baseline excitatory weight is invariant across development in our simulations.

To start the dev-STP simulation, we used the baseline STD parameters given in Table 2 at the beginning of the simulation, and slowly changed the parameters from depressing to facilitating at excitatory synapses. Toward this end, we averaged the postsynaptic neuron’s firing rate over a 500ms window and monitored how often it exceeded the ISP target rate of 5Hz by way of a variable *x*_exceed_ that was updated as follows

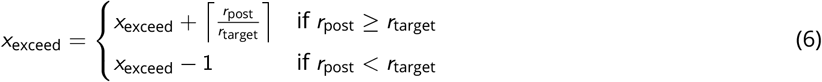

where *r*_post_ is the postsynaptic firing rate and *r*_target_ is the ISP target rate (see above). We increment STP to the next set of more facilitating STP parameters when *x*_exceed_ ≤ 0. In other words, the STP parameters are incremented only when the postsynaptic firing rate is equal to or below the ISP target rate for a sufficient period of time, i.e. a time that is proportional to the degree to which the postsynaptic firing rate has exceeded the target rate in the recent past. Changing the excitatory STP to a more facilitating state raises the postsynaptic firing rate, which increases *x*_exceed_, thus preventing further facilitating changes in STP until inhibitory synaptic weights strengthen and subsequently decrease the postsynaptic firing rate to the target rate, and the cycle starts over. Eventually, the STP parameter values reach the final (experimentally observed ^8^) STF parameter values (given in Table 2).

For both the fixed-STF and fixed-STD simulations, STP parameters at all excitatory synapses were set to depression and facilitation (Table 2), respectively, for the duration of the simulation.

Further, we quantified the level of “pathological activity” in all three models as the cumulative difference between the observed firing rate and the target firing rate for all input channels (Fig. 2g.i). We also considered the variability of firing rates, i.e. the coefficient of variation (standard deviation divided by the mean) of the firing rates averaged across 10s bins using a sliding window (Fig. 2g.ii).

#### Variants of developmental STP model

We conducted additional simulations to test three variants of the dev-STP model introduced above. In the first control variant we normalized the steady-state PSP amplitudes when using a 5 Hz presynatic Poisson input (Fig. S2) instead of normalizing to the first PSP. STP parameters in this dev-STP model were modified over development as described above. In this variant, the fixed-STF model displayed a lower initial firing rate than that of the standard model (Fig. S2b), failing to reach the ISP target rate and experimentally observed firing rates in young animals ^27–30^. Receptive field development in this variant is otherwise qualitatively similar to our dev-STP model, if somewhat more slowly (Fig. S2g).

In the second control variant we normalized the steady-state PSP of both STD and STF to be equal when using a 10Hz (instead of 5Hz as in the standard model) presynatic Poisson input (Fig. S3). In this case, STF was weakened enough that fixed-STF in young animals exhibited firing rates near the ISP target rate as observed experimentally ^27–30^. However, because of weakened STF, the model failed to develop fine-tuned tuning curves over development (Fig. S3f-h).

Finally, we tested a variant of our model in which the developmental shift from STD in young neurons to STF in adult neurons was not activity-dependent. Instead, we altered the dev-STP model to a model in which STP changes occurred at fixed intervals of 3 seconds (Fig. S4e). If these changes occur too frequently, unstable dynamics unfolded so some fine tuning of how often STP changes was required. This third variant also produced qualitatively similar results to our standard dev-STP model (compare Fig. S4, Fig. 2).

#### Excitatory and inhibitory tuning curves

To calculate the excitatory and inhibitory tuning curves, we monitored the excitatory and inhibitory conductances for each of the 8 input channels separately, and calculated the respective currents using

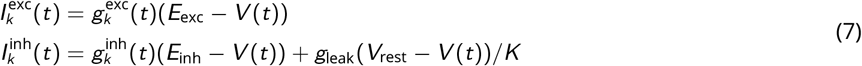

where 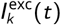 and 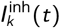 are the excitatory and inhibitory currents and 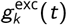 and 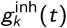 are the excitatory and inhibitory conductances of the *k*-th channel at time *t*, respectively ^21^. *E*_exc_ and *E*_inh_ are the excitatory and inhibitory reversal potentials, respectively. *V* (*t*) is the postsynaptic membrane potential at time *t*, *g*_leak_ is the leaky conductance, and *V*_rest_ is the resting membrane potential. After calculating the excitatory and inhibitory currents for each channel at all time points, we averaged the excitatory and inhibitory currents across 10 seconds to generate the tuning curves shown in the figures.

#### Output response dynamics across development

To measure how the neuron output response changed over the course of simulated development, we stopped the dev-STP simulation (Fig. 2) at 10s, 500s, 1,000s, 2,000s, 10,000s, and 30,000s simulated time and examined the neuron response dynamics of the model. For each snapshot, we ran 50 step current trials with frozen parameters and compared the average firing rates of the dev-STP scenario with those of the fixed-STD and fixed-STF scenarios (Fig. 4b).

To investigate how input tuning changed over development, we calculated the cross correlations between the input and output rates for each of the 8 channels ^21^. We obtained the correlation range by subtracting the minimum from the maximum correlation and normalized the range by dividing by the mean correlation of all channels with the output (Fig. 4d).

#### Signal transmission across development

To investigate signal transmission across development, we presented a 250ms long 150Hz input stimulus to the preferred input channel every 100 seconds of the dev-STP simulation (Fig. 2). We analysed the output firing rates during the first 50ms after stimulus onset (phasic period) and the remaining 200ms afterwards (tonic period); Fig. 5b,c). We also tested a double step input stimulus, two 250ms 150 Hz input stimuli separated by 250ms of spontaneous activity (Fig. 5d,e).

#### Temporal precision simulations

We compared the temporal precision of postsynaptic spikes in our model with experimental observations ^25^. To this end, we stimulated the preferred channel (5) of the output neuron with a 200ms step current, imitating a pure tone in the preferred frequency in the auditory cortex ^25^. To quantify the temporal precision of the response, we calculated the standard deviation of the delay between the stimulus onset and the first postsynaptic spike, denoted as the jitter ^25^. To allow comparison across different firing rates, we also calculated a normalized jitter, i.e., the jitter’s coefficient of variation. The normalized jitter was compared for different preferred-channel stimulus strengths as well as for varying spontaneous activity levels (Fig. 6c-e).

#### Working memory

To test for working memory-like properties, we used two simulation protocols. In the “memory preloaded” trials, we stimulated the neuron with a 300ms long 150Hz steady state stimulus (a memory) in the preferred channel. All remaining channels received spontaneous rates at 5Hz. After the memory preloading period, the preferred channel input received spontaneous firing rate inputs for a 300ms delay period, followed by a weaker, 100ms long 50Hz “recall cue” stimulus. For “control” trials, the input channels of the neuron only received the 100ms recall cue, to the preferred channel, without preloading.

We then compared the firing rates during recall between the memory preloaded and control trials, to study the ‘silent’ working memory effects in our model. We tested this throughout simulated development, by freezing the dev-STP simulation every 50s and simulating 500 trials of the memory-preloaded simulations and 500 trials of the control simulations.

#### Simulator

Simulations were conducted in Python using Brian Simulator 2. Code to reproduce our key findings is available at github.com/djia/dev-stp.

---

**Figure S1.**
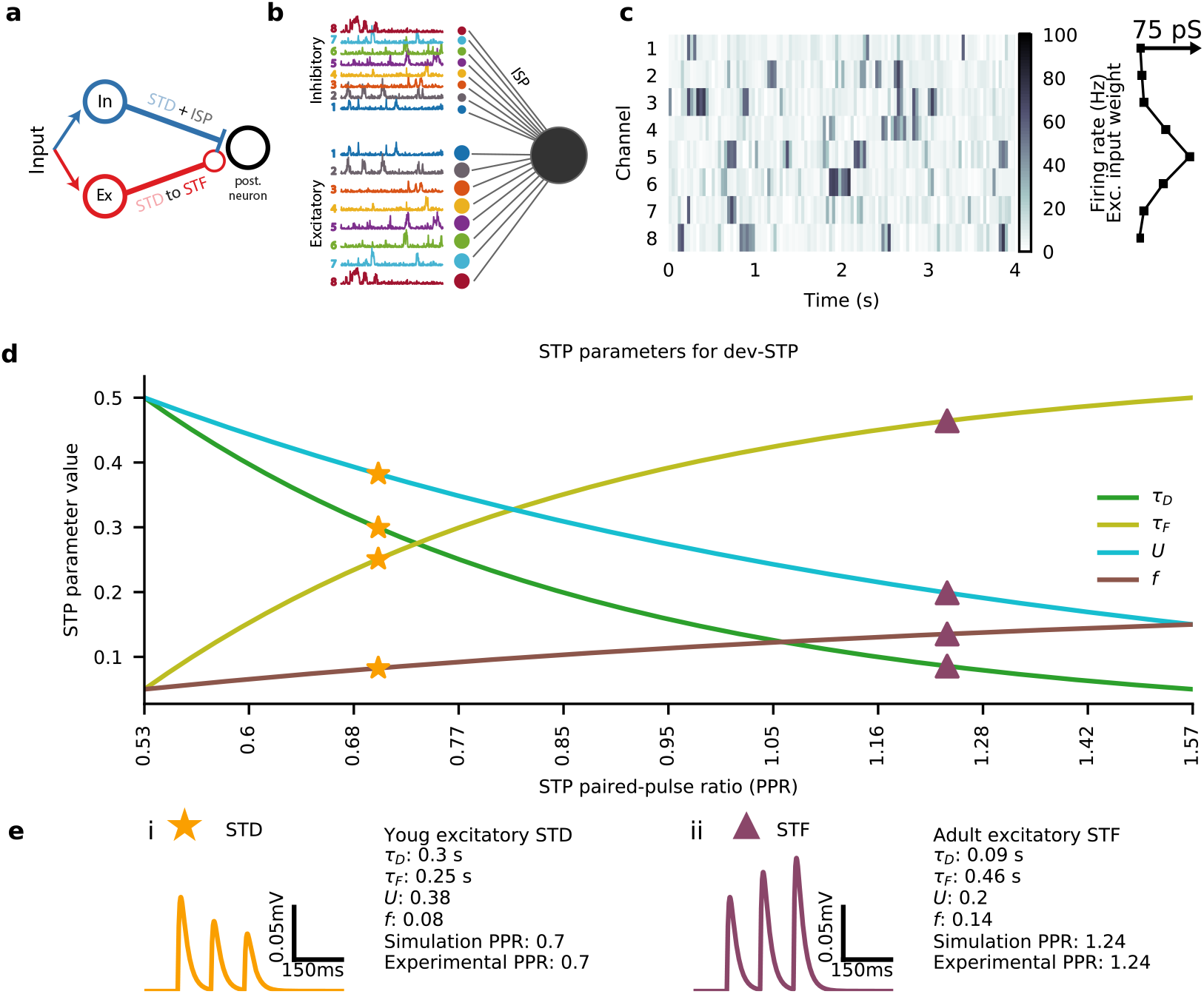
Details of the cortical circuit and plasticity models. **(a)** Schematic of a single channel feedforward circuit with correlated excitatory and inhibitory input, and the respective forms of plasticity. **(b)** Feedforward neural circuit with 8 channels and correlated excitatory and inhibitory inputs. **(c)** Left: example of input given to the 8 channel feedforward neural circuit; right: excitatory tuning curve strength for each of the 8 channels. **(d)** Each of the four STP parameters, *τ_D_*, *τ_F_*, *U*, and *f* resulting in different paired-pulse ratios (PPRs) (Table 2). Parameters matching the young (orange star) and adult (purple triangle) STP PPRs as used in the dev-STP model are highlighted. **(e)** Example postsynaptic potential traces for the STP parameter values of both young (i) and adult animals (ii; cf. **d**).

**Figure S2.**
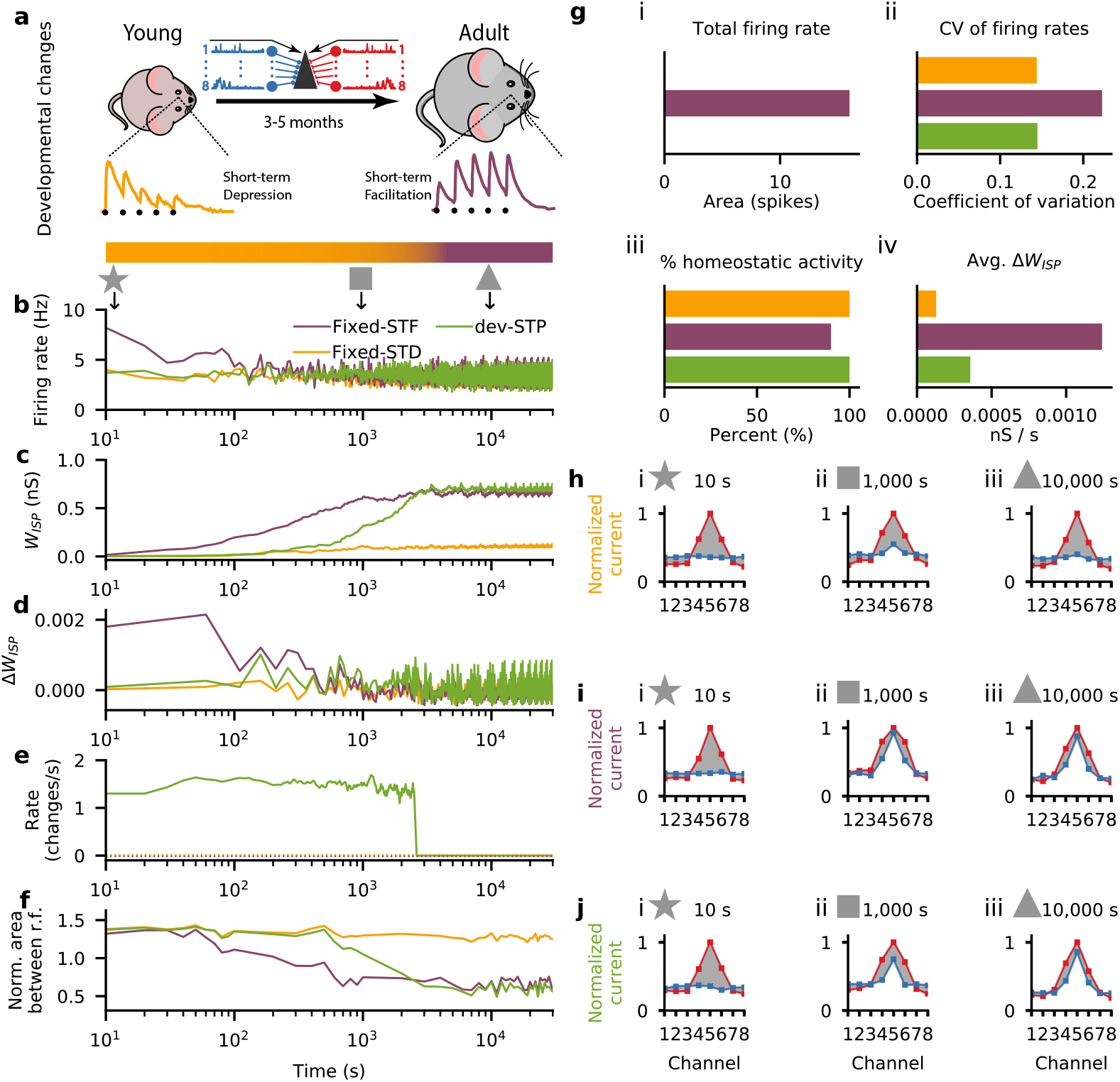
Developmental STP model with depression and facilitation normalized to the steady-state firing rate at 5Hz input. (**a**) Schematic of our developmental short-term plasticity (STP) model (cf. Fig. S1); top: young and adult STP (as in Fig. 1); bottom: gradual changes in STP from depressing to facilitating dynamics (orange and purple respectively, in log-scale as in b-f). (**b**-**f**) Different variables of the model across simulated development for three different models: fixed short-term depression (fixed-STD, orange), fixed short-term facilitation (fixed-STF, purple) and developmental model with gradual changes in STP (dev-STP, green line). Note x-axis on log-scale. (**b**) Receiver neuron firing rate. (**c**) Mean inhibitory weight. (**d**) Mean changes in the weight of the inhibitory synaptic afferents. (**e**) Rate of STP change (note that both fixed-STF and STD remain fixed, shown as dashed lines). (**f**) Area between normalised excitatory and inhibitory tuning curves (cf. h-j) during the course of simulated development. A normalised area close to 0 represents a perfectly balanced neuron. (**g**) Additional statistics for the three models. (i) Total neuronal activity calculated using the area between the firing rate in (b) and the desired target rate of 5 Hz. (ii) Average coefficient of variation of the firing rates across simulated development (cf. (b)). (iii) Percent of time spent under homeostasis (i.e. at the desired firing rate; cf. (b)). (iv) Average change in inhibitory weights (cf. (d)). (**h**-**j**) Snapshots of excitatory and inhibitory tuning curves across three points in simulated development: 10s (star), 1000s (square) and 10 000s (triangle). Shaded gray area represents difference between excitatory and inhibitory tuning curves (cf. (f)). (**h**-**j**) Excitatory (red) and inhibitory (blue) postsynaptic tuning curve for the fixed-STD (h), fixed-STF (i) and dev-STP models (j).

**Figure S3.**
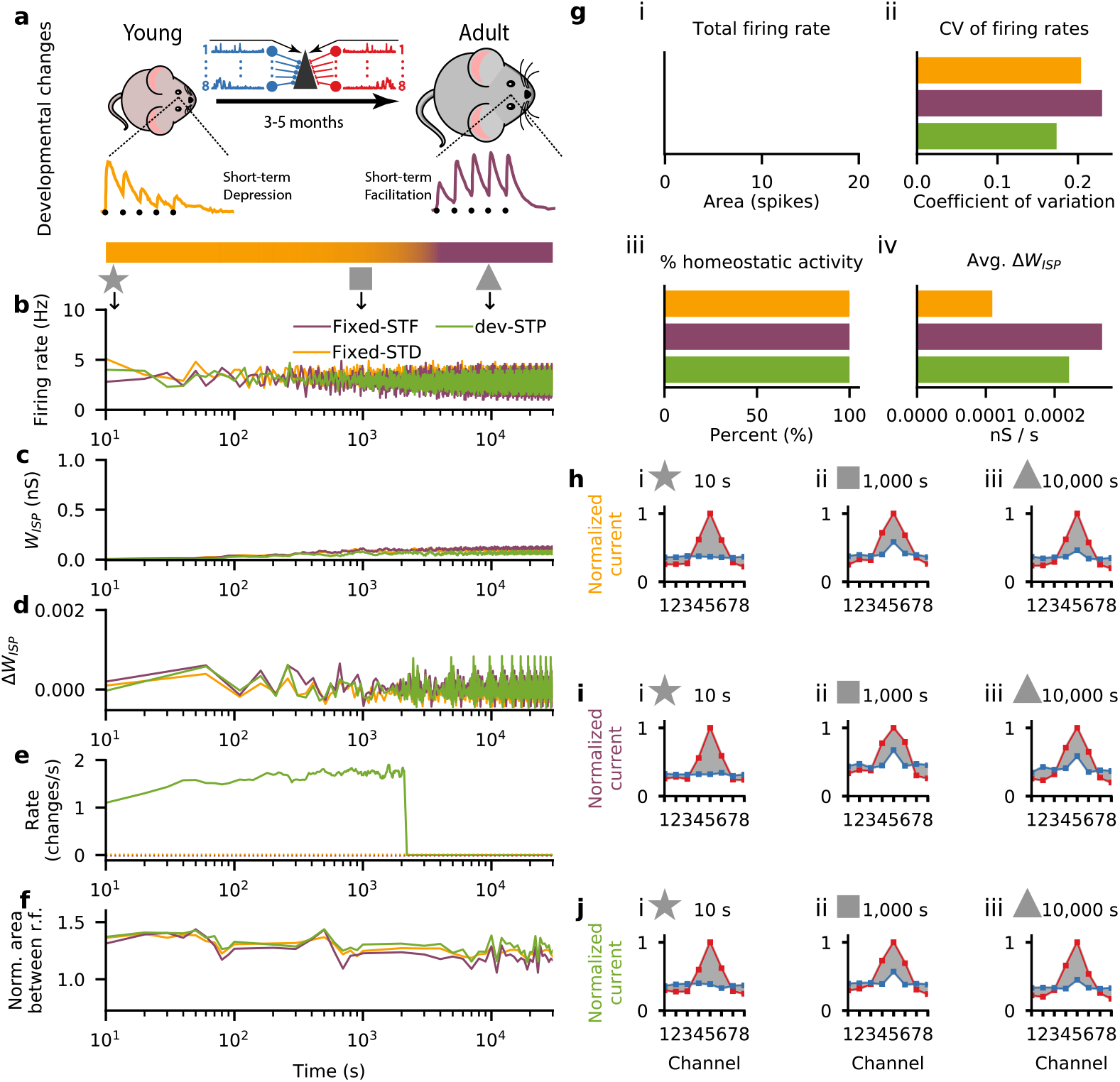
Developmental STP model with depression and facilitation normalized to the steady-state firing rate at 10Hz input. (**a**) Schematic of our developmental short-term plasticity (STP) model (cf. Fig. S1); top: young and adult STP (as in Fig. 1); bottom: gradual changes in STP from depressing to facilitating dynamics (orange and purple respectively, in log-scale as in b-f). (**b**-**f**) Different variables of the model across simulated development for three different models: fixed short-term depression (fixed-STD, orange), fixed short-term facilitation (fixed-STF, purple) and developmental model with gradual changes in STP (dev-STP, green line). Note x-axis on log-scale. (**b**) Receiver neuron firing rate. (**c**) Mean inhibitory weight. (**d**) Mean changes in the weight of the inhibitory synaptic afferents. (**e**) Rate of STP change (note that both fixed-STF and STD remain fixed, shown as dashed lines). (**f**) Area between normalised excitatory and inhibitory tuning curves (cf. h-j) during the course of simulated development. A normalised area close to 0 represents a perfectly balanced neuron. (**g**) Additional statistics for the three models. (i) Total neuronal activity calculated using the area between the firing rate in (b) and the desired target rate of 5 Hz. (ii) Average coefficient of variation of the firing rates across simulated development (cf. (b)). (iii) Percent of time spent under homeostasis (i.e. at the desired firing rate; cf. (b)). (iv) Average change in inhibitory weights (cf. (d)). (**h**-**j**) Snapshots of excitatory and inhibitory tuning curves across three points in simulated development: 10s (star), 1000s (square) and 10 000s (triangle). Shaded gray area represents difference between excitatory and inhibitory tuning curves (cf. (f)). (**h**-**j**) Excitatory (red) and inhibitory (blue) postsynaptic tuning curve for the fixed-STD (h), fixed-STF (i) and dev-STP models (j).

**Figure S4.**
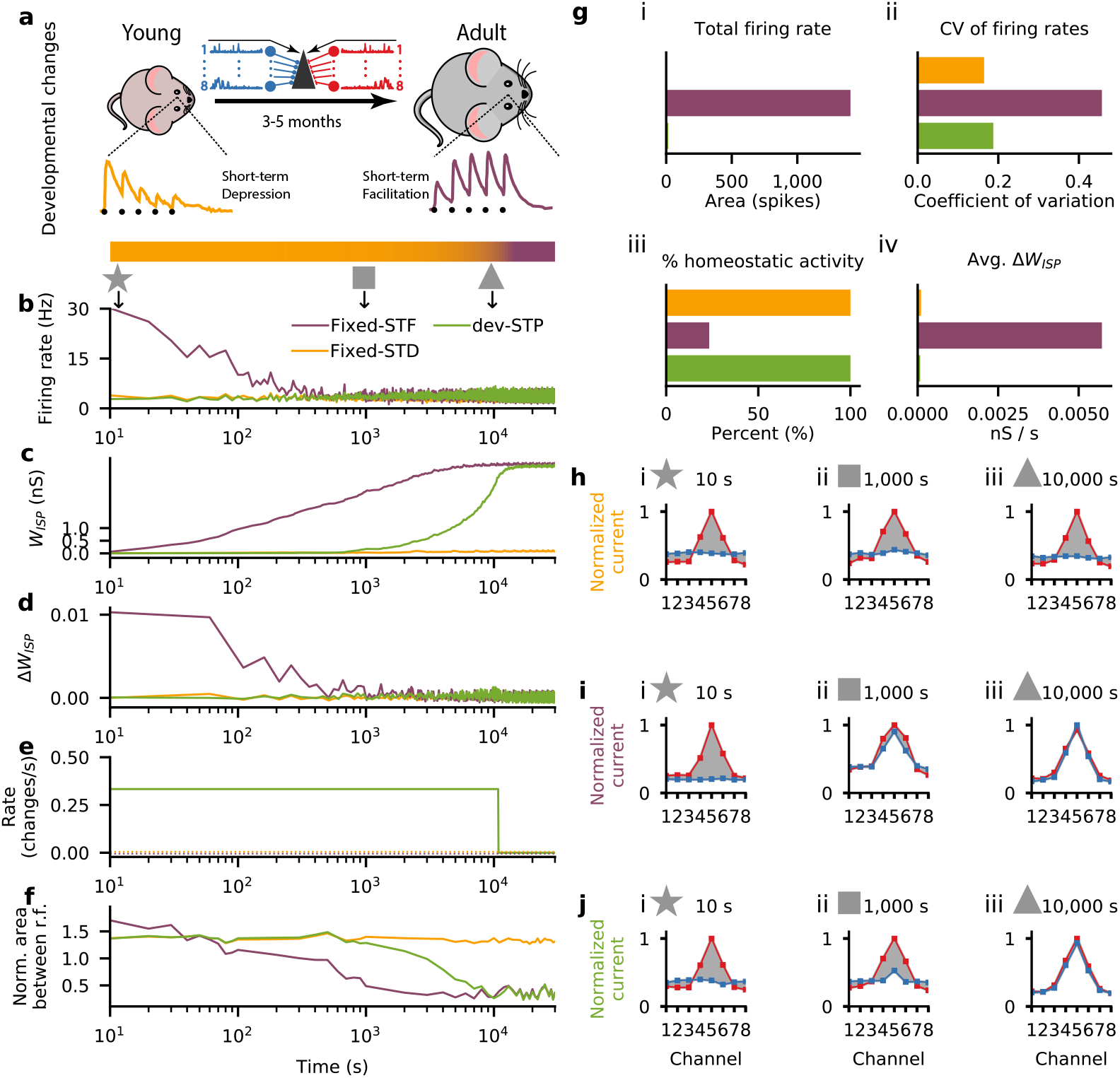
Developmental STP model in which STP changes are pre-defined. (**a**) Schematic of our developmental short-term plasticity (STP) model (cf. Fig. S1); top: young and adult STP (as in Fig. 1); bottom: gradual changes in STP from depressing to facilitating dynamics (orange and purple respectively, in log-scale as in b-f). (**b**-**f**) Different variables of the model across simulated development for three different models: fixed short-term depression (fixed-STD, orange), fixed short-term facilitation (fixed-STF, purple) and developmental model with gradual changes in STP (dev-STP, green line). Note x-axis on log-scale. (**b**) Receiver neuron firing rate. (**c**) Mean inhibitory weight. (**d**) Mean changes in the weight of the inhibitory synaptic afferents. (**e**) Rate of STP change (note that both fixed-STF and STD remain fixed, shown as dashed lines). (**f**) Area between normalised excitatory and inhibitory tuning curves (cf. h-j) during the course of simulated development. A normalised area close to 0 represents a perfectly balanced neuron. (**g**) Additional statistics for the three models. (i) Total neuronal activity calculated using the area between the firing rate in (b) and the desired target rate of 5 Hz. (ii) Average coefficient of variation of the firing rates across simulated development (cf. (b)). (iii) Percent of time spent under homeostasis (i.e. at the desired firing rate; cf. (b)). (iv) Average change in inhibitory weights (cf. (d)). (**h**-**j**) Snapshots of excitatory and inhibitory tuning curves across three points in simulated development: 10s (star), 1000s (square) and 10 000s (triangle). Shaded gray area represents difference between excitatory and inhibitory tuning curves (cf. (f)). (**h**-**j**) Excitatory (red) and inhibitory (blue) postsynaptic tuning curve for the fixed-STD (h), fixed-STF (i) and dev-STP models (j).

**Figure S5.**
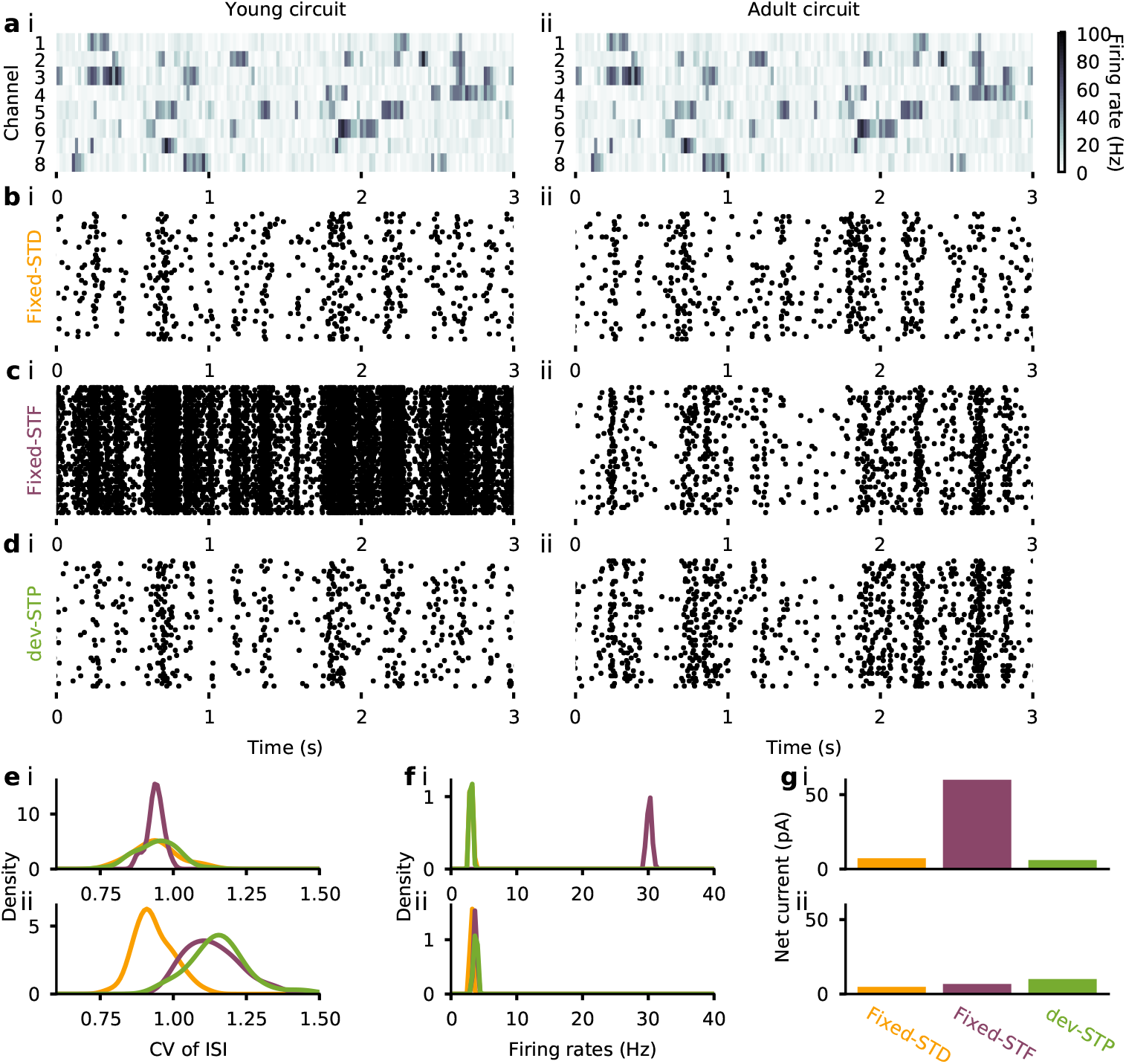
Developmental STP shapes firing statistics. (**a**) Input activity for each of the 8 channels over 3 seconds. Activity at the start of simulated development (i, young condition) and after 8 hours of simulation (ii, adult condition) as in Fig. 2; color code represents firing rate of input. (**b-d**) Raster plot of receiver neuron for fixed-STD model (b), fixed-STF (c) and developmental STP model (d). (**e-g**) Summary statistics of the three models (as in b-d) for both young (i) and adult conditions (ii). (**e**) Coefficient of variation of the inter-spike intervals. (**f**) Average firing rates of the receiver neuron over 50 trials. (**g**) Average net current of the receiver neuron.

**Figure S6.**
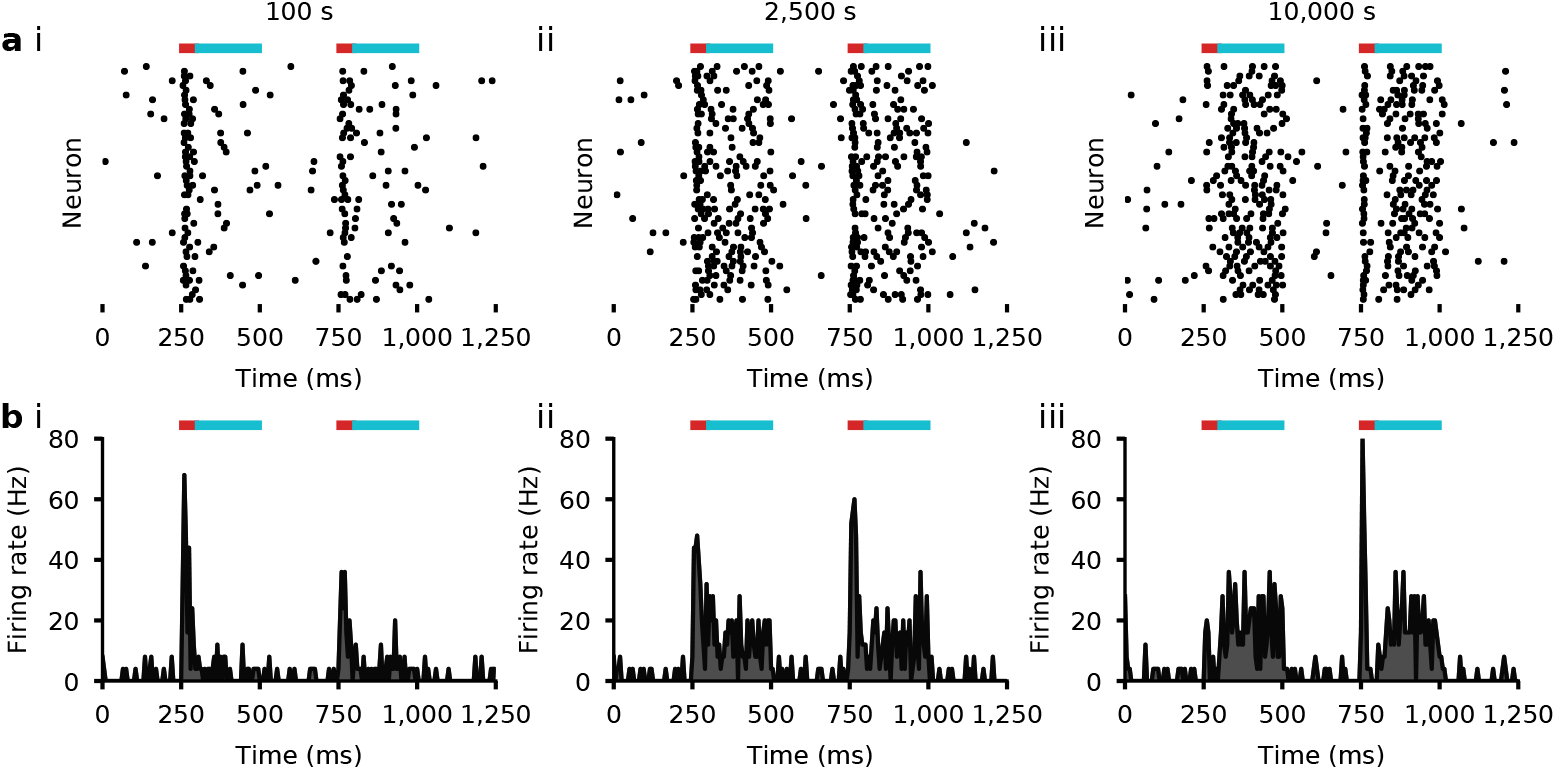
Spike rasters for step inputs at various snapshots. (**a**) Spike responses to two 150Hz step inputs to the preferred channel when using the dev-STP model at 100s (i), 2,500s (ii), and 10,000s (iii); color bars on top represent the time at which the step inputs were given; the first 50ms corresponds to the phasic activity (red), and the rest of the input time period to the tonic activity (cyan); **b** Firing rates of the spikes in (a) averaged across trials using 5ms bins.

